# Prefrontal D1 dopamine-receptor neurons and delta resonance in interval timing

**DOI:** 10.1101/216473

**Authors:** Young-Cho Kim, Nandakumar S. Narayanan

## Abstract

Considerable evidence has shown that prefrontal neurons expressing D1-type dopamine receptors (D1DRs) are critical for working memory, flexibility, and timing. This line of work predicts that frontal neurons expressing D1DRs mediate cognitive processing. During timing tasks, one form this cognitive processing might take is time-dependent ramping activity — monotonic changes in firing rate over time. Thus, we hypothesized the prefrontal D1DR+ neurons would strongly exhibited time-dependent ramping during interval timing. We tested this idea using an interval-timing task in which we used optogenetics to tag D1DR+ neurons in the mouse medial frontal cortex (MFC). While 23% of MFC D1DR+ neurons exhibited ramping, this was significantly less than untagged MFC D1DR+ neurons. By contrast, MFC D1DR+ neurons had strong delta-frequency (1-4 Hz) coherence with other MFC ramping neurons. This coherence was phase-locked to cue onset and was strongest early in the interval. To test the significance of these interactions, we optogenetically stimulated MFC D1DR+ neurons early vs. late in the interval. We found that 2-Hz stimulation early in the interval was particularly effective in rescuing timing-related behavioral performance deficits in dopamine-depleted animals. These findings provide insight into MFC networks and have relevance for disorders such as Parkinson’s disease and schizophrenia.

**Significance Statement:** Prefrontal D1DRs are involved in cognitive processing and cognitive dysfunction in human diseases such as Parkinson’s disease and schizophrenia. We use optogenetics to identify these neurons, as well as neurons that are putatively connected to MFC D1DR+ neurons. We study these neurons in detail during an elementary cognitive task. These data could have relevance for cognitive deficits for Parkinson’s disease, schizophrenia, and other diseases involving frontal dopamine.

## Introduction

Medial and lateral regions of the mammalian frontal cortex are involved in cognitive processes such as working memory, flexibility, and timing (Fuster 2008). Frontal neurons encode intricacies of cognitive processing such as remembered items, prospective action, errors, goals, and temporal control of action (Niki and Watanabe 1979; Goldman-Rakic et al. 2004; Miller and D’Esposito 2005; Narayanan, Cavanagh, et al. 2013; Ma et al. 2014; Hardung et al. 2017). The coordinated activity of these neurons is detectable by macro-level techniques such as functional magnetic resonance imaging (fMRI) or electroencephalography (EEG). For instance, during tasks requiring cognitive control, frontal EEG electrodes often detect low-frequency oscillations in delta (1-4 Hz) and theta (4-8 Hz) frequency bands (Cavanagh and Frank 2014; Parker, Chen, et al. 2015; Parker et al. 2017a; Chen et al. 2016; Kim et al. 2017).

Approximately 20% of frontal cortical neurons express D1-type dopamine receptors (D1DRs; Gaspar et al. 1995). In humans, PET imaging studies have implicated D1DRs in working memory (Okubo et al. 1997; Abi-Dargham et al. 2002). In primates and rodents, local infusions of drugs targeting D1DRs impair performance on working memory, inhibitory control, flexibility, and timing tasks (Sawaguchi and Goldman-Rakic 1991, 1994; Vijayraghavan et al. 2007; St Onge et al. 2011; Parker, Alberico, et al. 2013; Parker, Ruggiero, et al. 2015; Jenni et al. 2017). D1DR agonists and antagonists can specifically attenuate neuronal activity related to working memory (Williams and Goldman-Rakic 1995; Vijayraghavan et al. 2007) and temporal processing (Narayanan et al. 2012a; Parker, Andreasen, et al. 2013; Parker, Chen, et al. 2014, 2015; Parker, Ruggiero, et al. 2015; Narayanan 2016). During interval-timing tasks, this temporal processing can take the form of *time-related ramping*, which involves monotonic increases or decreases in neuronal firing rate across temporal intervals. Drugs acting on D1DRs specifically attenuate time-related ramping during interval-timing tasks (Parker, Chen, et al. 2014; Parker et al. 2017a; Parker, Ruggiero, et al. 2015). Finally, stimulating MFC D1DR+ neurons can increase ramping activity and improve performance of interval-timing tasks (Narayanan et al. 2012a; Kim et al. 2017). These data lead to the specific hypothesis that MFC D1DR+ neurons strongly exhibit time-related ramping activity.

We tested this hypothesis by using optogenetics to tag putative MFC D1DR+ neurons in D1-Cre mice during interval-timing tasks. To our surprise, we did not find evidence to support the hypothesis that a large fraction of MFC D1DR+ neurons exhibited ramping activity. Instead, our data suggest that MFC D1DR+ neurons had delta/theta coherence with other MFC ramping neurons. Our findings could have relevance for our fundamental understanding of cortical networks and for human diseases involving impaired frontal dopamine.

## Results

We trained 6 mice to perform a fixed-interval task with a 12-s interval (Fig 1A). In these animals, we implanted optrodes targeting the MFC (Fig 1B). We isolated 314 MFC neurons. We used optogenetic tagging to identify MFC D1DR+ neurons by recording from D1-Cre mice virally expressing AAV-DIO-ChR2 in the MFC, which expresses ChR2 in MFC D1DR+ neurons. In recording experiments, MFC neurons that spiked within <5 ms of 473-nm laser onset were considered putative MFC D1DR+ neurons (Fig 2). We identified 93 tagged MFC D1DR+ neurons. Optogenetic stimulation did not affect waveform shape (average correlation coefficient with non-stimulated spikes: r=0.99). The average latency of tagged neurons was 1.99 ms and the jitter was 0.82 ms.

**Figure 1:**
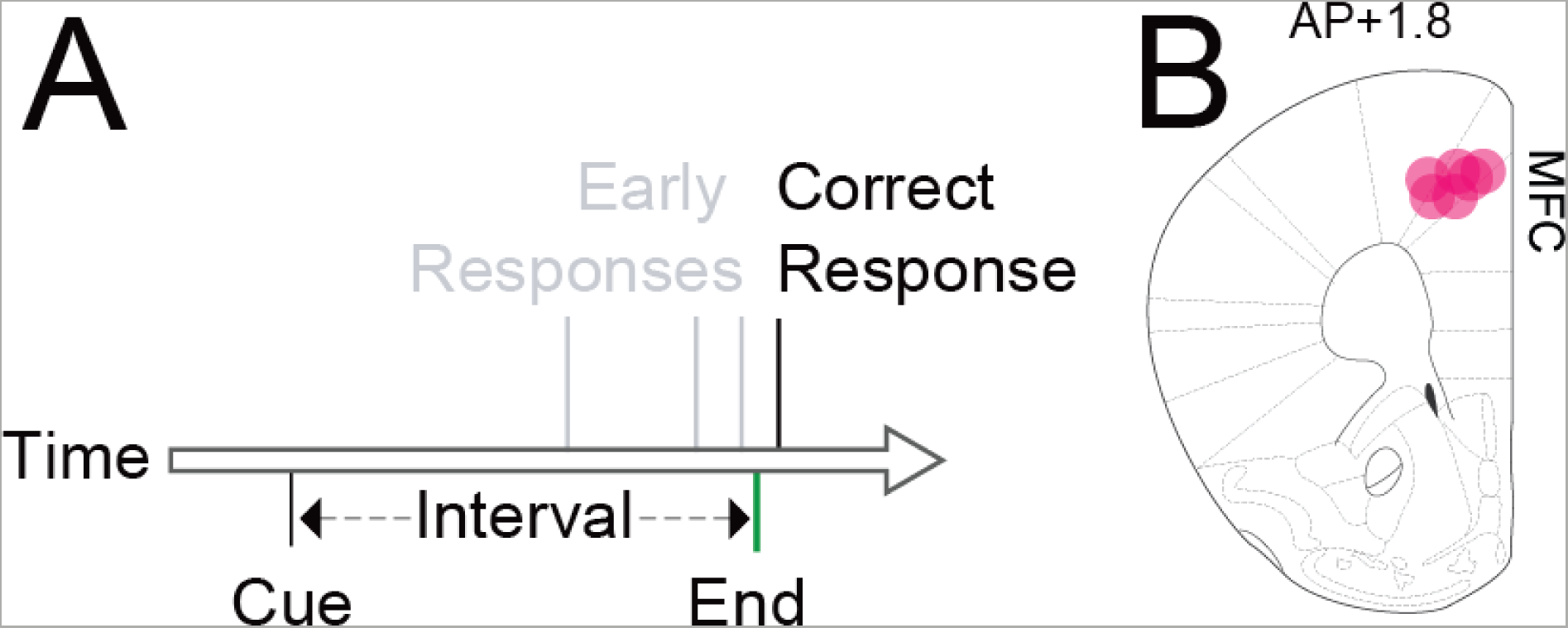
Interval timing requires mesocortical dopamine. A) We trained mice to perform a fixed-interval timing task, in which rodents had to make a motor response 12 s after a starting cue. Early responses were not reinforced, whereas the first response after interval end was rewarded. B) We recorded from 6 mice with optrodes placed in medial frontal cortex (MFC; red dots).

**Figure 2:**
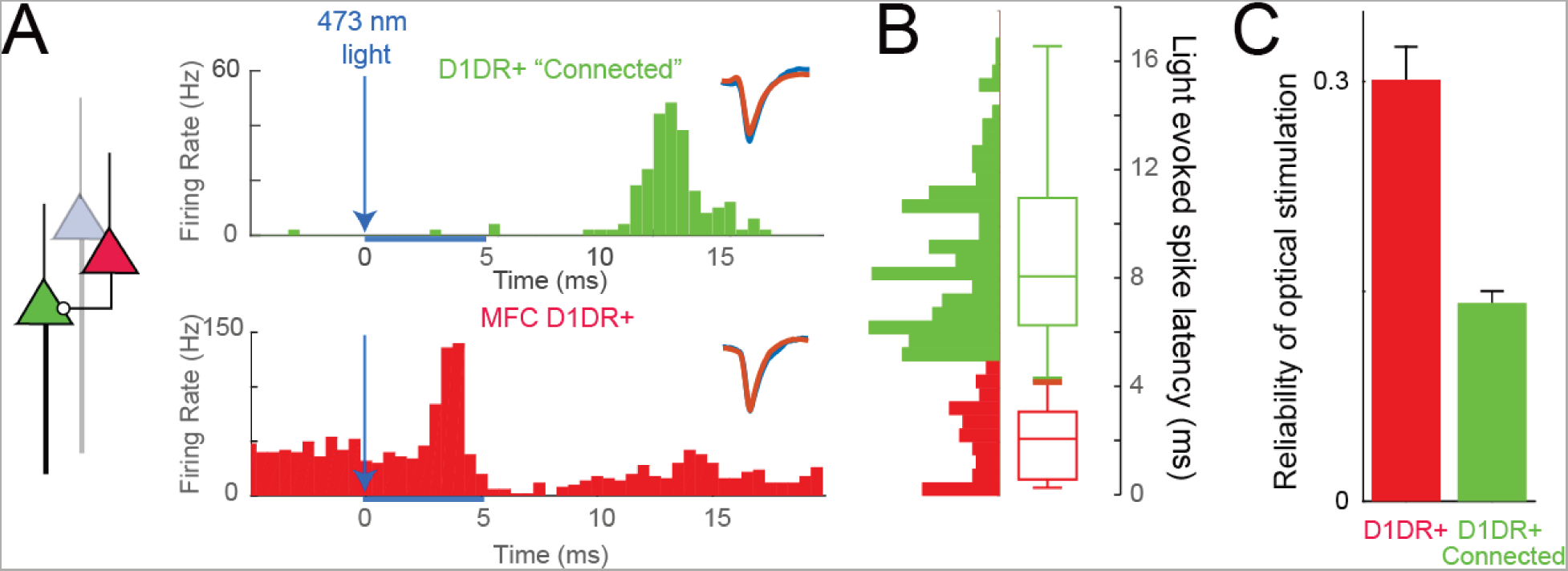
Optogenetic tagging of MFC D1DR+ and connected neurons. A) By virally expressing DIO-ChR2 in the MFC of D1-Cre mice, we can optogenetically ‘tag’ MFC D1DR+ neurons (in red) as those with short-latency action potentials <5 ms from 473-nm laser onset, as this latency is most consistent with action potentials initiated within the recorded neuron rather than synaptic transmission. Aside from tagged neurons however, we noticed a second peak in latency in some neurons that could be consistent with monosynaptic connectivity to MFC D1DR+ neurons. We operationally termed neurons with this peak MFC D1DR+ ‘Connected’ neurons (in green). B) Light-evoked latencies as histograms and box plots of putative MFC D1DR+ (red) and putative MFC D1DR+ Connected neurons are shown. Waveforms for non-stimulation (orange) and stimulated trials (blue) inset. C) The reliability of action potentials triggered by optical stimulation pulse is significantly higher in D1DR+ neurons than D1DR+ connected neurons.; * −p<0.05.

We also noticed that some MFC neurons had clear peaks in firing rate 6 – 18 ms after laser stimulation (Fig 2B). We operationally described these neurons as putatively connected with MFC D1DR+ neurons and termed them ‘D1DR+ Connected’ neurons. 149 MFC neurons met these criteria and were putatively labeled as D1DR+ Connected. Laser-induced spikes were more reliable in tagged D1DR+ neurons than in D1DR+ Connected neurons (0.30 ± 0.02 vs 0.14 ± 0.008, t_(471)_=8.003, p<0.0001; Fig 2C). Because of limitations in our recording techniques, it is difficult to make further conclusions about the anatomical or synaptic configuration of these neurons. In the remaining 72 neurons, we could identify no discernable change in spike rate with optogenetic stimulation. These neurons were labeled as ‘Untagged’. D1DR+ neurons had similar firing rates to D1DR+ Connected and untagged neurons (D1DR+: 8.4 ± 1.2 Hz; D1DR+ Connected: 7.9 ± 0.9 Hz; Untagged: 9.8 ± 1.3 Hz).

We explored how MFC D1DR+ neurons, MFC D1DR+ Connected neurons, and untagged MFC neurons were involved in interval timing. Past work by our group and others (Narayanan et al. 2012b; Kim et al. 2013, 2017; Xu et al. 2014; Gouvea et al. 2015; Parker, Ruggiero, et al. 2015; Emmons et al. 2017) has identified three common patterns of MFC activity: stimulus-related activity (Fig 3A), temporal processing in the form of time-related ramping activity (Fig 3B), or response-related activity (Fig 3C). Time-related ramping was defined as a monotonic increase or decrease in firing rate as quantified by linear regression of firing rate over the interval (Narayanan 2016; Kim et al. 2017). Past studies have demonstrated that MFC D1DR pharmacological or optogenetic manipulation can selectively influence ramping leading to our hypothesis that MFC D1DR+ neurons ramp (Parker, Chen, et al. 2014; Parker, Ruggiero, et al. 2015; Kim et al. 2017).

**Figure 3:**
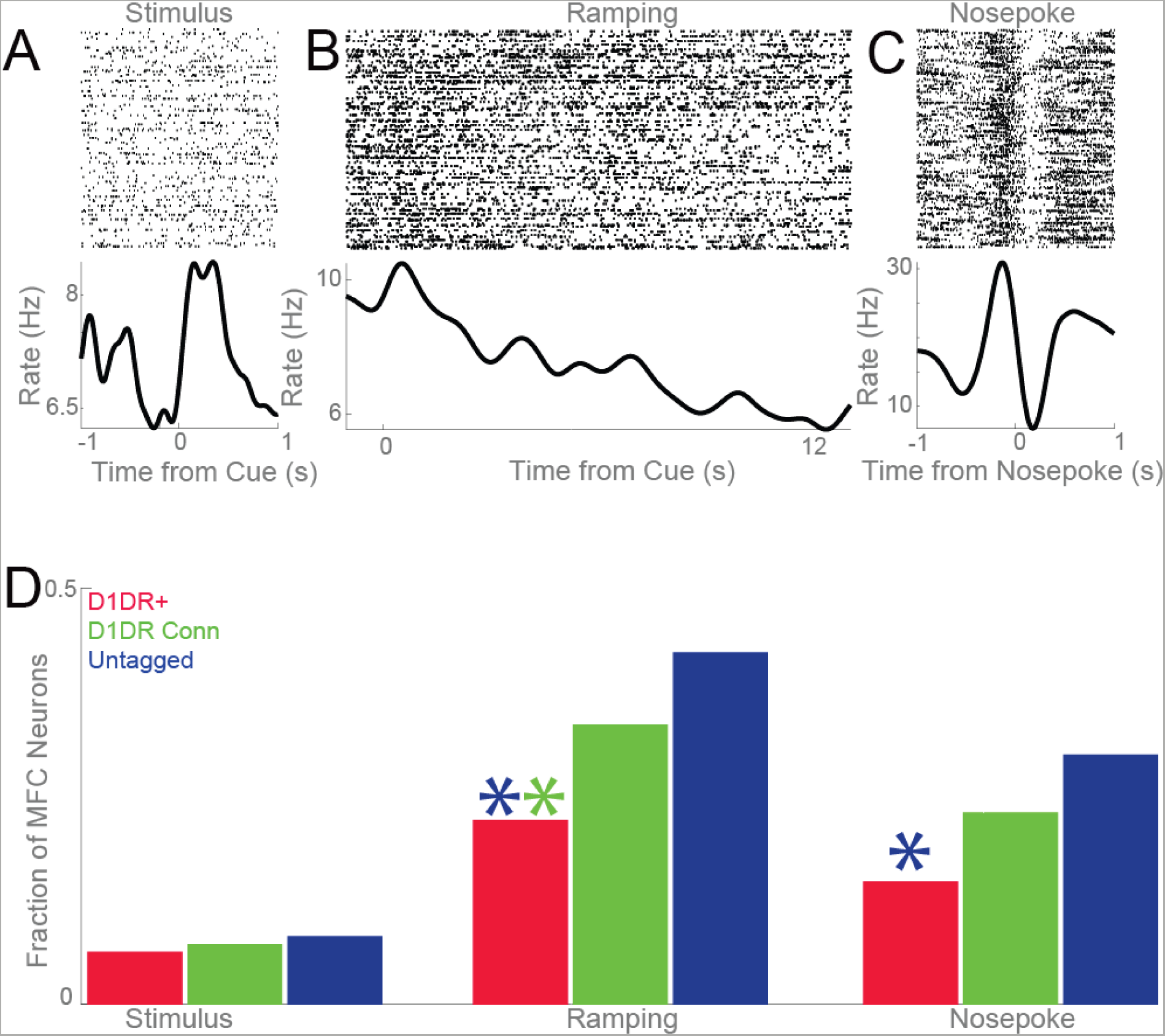
MFC D1DR+ neurons have less time-related ramping than untagged MFC neurons. Examples of A) a stimulus-modulated neuron, B) a neuron with time-related ramping, and C) a nosepoke-modulated neuron. D) We examined the modulation patterns of MFC D1DR+ neurons (red), MFC D1DR+ Connected neurons (green), and untagged MFC neurons (blue). There were less MFC D1DR+ ramping neurons than MFC D1DR+ Connected (green asterisk) and untagged MFC neurons (blue asterisk). There were also less MFC D1DR+ than untagged MFC nosepoke neurons (blue asterisk). Data from 314 MFC neurons in 6 mice, including 93 MFC D1DR+ neurons. * indicates p<0.05.

We found that 21 of 93 MFC D1DR+ neurons (23%) exhibited time-related ramping. To our surprise, this was significantly less than MFC D1DR+ Connected neurons (51 of 149; 34%; χ^2^=3.7, p<0.05) and less than untagged MFC neurons (31 of 72, or 43%; χ^2^=8.9 p<0.003; Fig 3D). There were similar fractions of stimulus-related activity in MFC D1DR+ neurons and untagged neurons (Fig 3D). These data indicate that MFC D1DR+ neurons had *less* time-related ramping than other populations of MFC neurons. We also found that MFC D1DR+ neurons had less motor-related activity than untagged MFC neurons (χ^2^=4.8, p<0.03; Fig 3D), but not less than MFC D1DR+ Connected neurons.

To further analyze patterns of MFC neurons we turned to a data-driven method, principal component analysis (PCA). We have used PCA extensively in the past to capture patterns of neuronal activity (Chapin and Nicolelis 1999; Narayanan and Laubach 2009; Parker, Chen, et al. 2014; Emmons et al. 2017; Kim et al. 2017; Parker et al. 2017b). PCA revealed that the first four components each accounted for >10% of variance among MFC ensembles, with PC1 accounting for 22% of variance and PC4 accounting for 11% of variance (Fig 4A-C). Smaller components >PC5 accounted for progressively less variance and were not analyzed. In line with past work, PC1 exhibited time-related ramping (Narayanan and Laubach 2009; Narayanan 2016; Kim et al. 2017; Parker et al. 2017a). PC1 had significantly less loading on MFC D1DR+ neurons than untagged neurons (t_(163)_=3.6, p<0.0004; Fig 4C) or MFC D1DR+ Connected neurons (t_(240)_=2.1, p<0.03; Fig 4C). MFC D1DR+ neurons loaded strongly on PC4. This was stronger for MFC D1DR+ neurons than untagged neurons (t_(163)_=2.0, p<0.05) and MFC D1DR+ Connected neurons (t_(240)_=2.3, p<0.02). Taken together, these data do not support the hypothesis that MFC D1DR+ neurons exhibit ramping more strongly than MFC D1DR+ Connected or untagged MFC neurons.

**Figure 4:**
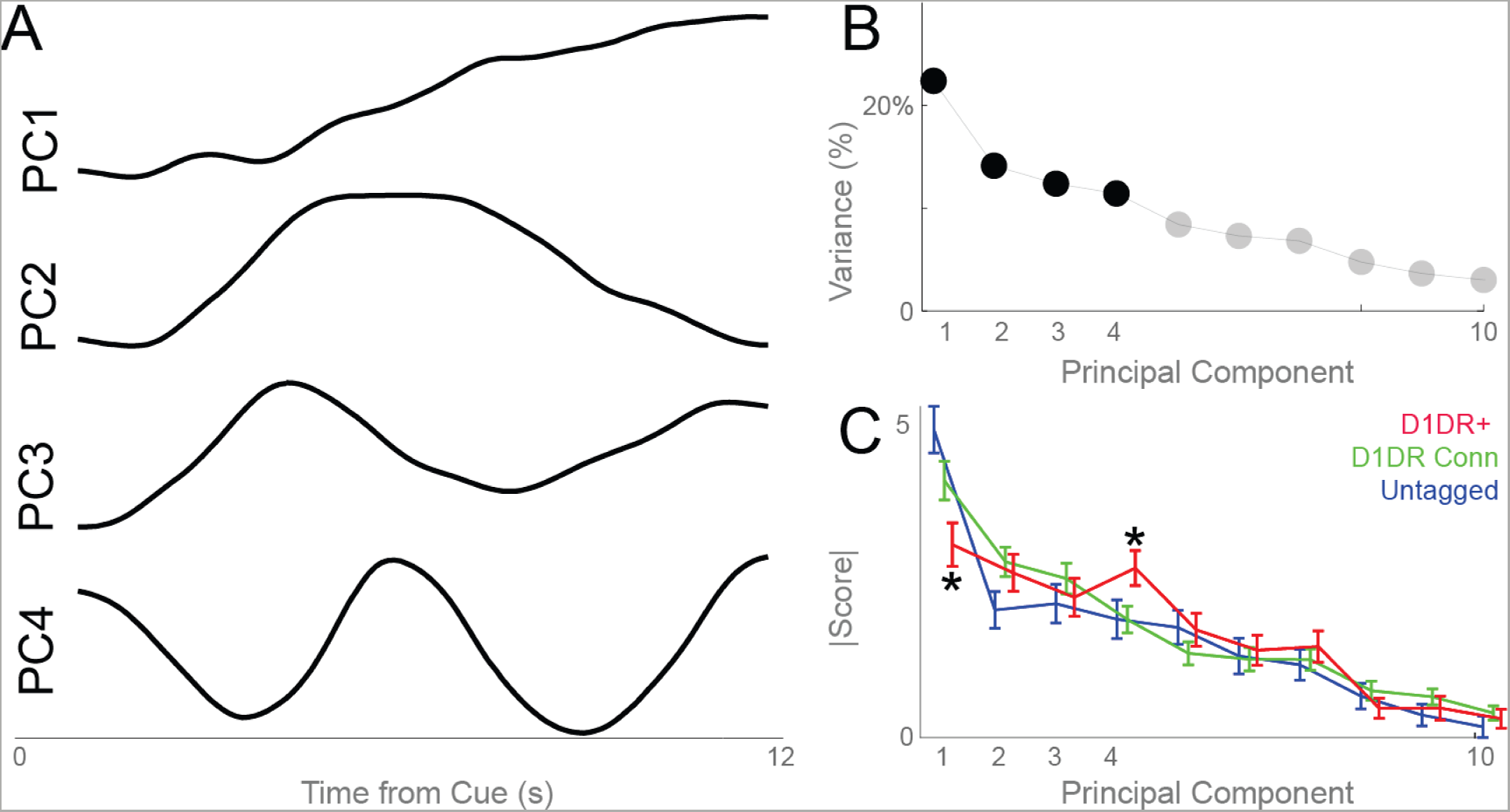
MFC D1DR+ neuronal ensembles have distinct principal components, including less ramping activity. Principal component analysis is a data-driven approach to identify dominant patterns in multivariate data. A) Four large components were noted; as in past work, PC1 was a ‘ramping’ component, and PC4 had an oscillatory pattern with a period of ~6 s. B) The first 4 PCs each explained >10% of variance. C) MFC D1DR+ neurons had less loading on PC1 and more loading on PC4. Data from 314 MFC neurons, including 93 MFC D1DR+ neurons, in 6 mice. * indicates p<0.05 for MFC D1DR+ vs. untagged MFC neurons.

Consequently, we investigated alternative patterns of MFC D1DR+ activity. Several prior studies by our group have indicated that low-frequency delta/theta rhythms around 4 Hz require MFC D1DRs (Cavanagh and Frank 2014; Parker, Narayanan, et al. 2014; Parker, Chen, et al. 2015; Parker, Ruggiero, et al. 2015; Chen et al. 2016). Accordingly, we examined spike-to-spike coherence between MFC D1DR+ neurons and D1DR+ Connected neurons as well as untagged neurons (Figure 5A). We found that MFC D1DR+ neurons had prominent cue-locked delta/theta coherence with ramping D1DR+ Connected neurons (Fig 5A). This was significantly more than we observed with non-ramping D1DR+ Connected neurons and untagged MFC neurons (Fig 5B-E; delta: t_(778)_=2.4, p<0.02; theta; t_(778)_=2.7, p<0.008; beta; t_(778)_=0.5, p<0.60). These data indicate that MFC D1DR+ neurons could have specific cue-triggered delta/theta synchrony with ramping neurons. We also noticed that this interaction was strongest early in the interval between 0 and 6 s at 1-8 Hz (Fig 5F; paired t_(250)_=2.9, p<0.004 first 6 s – last 6 s). These findings suggest that MFC D1DR+ neurons might interact with MFC ramping neurons via delta/theta interactions early in the interval.

**Figure 5:**
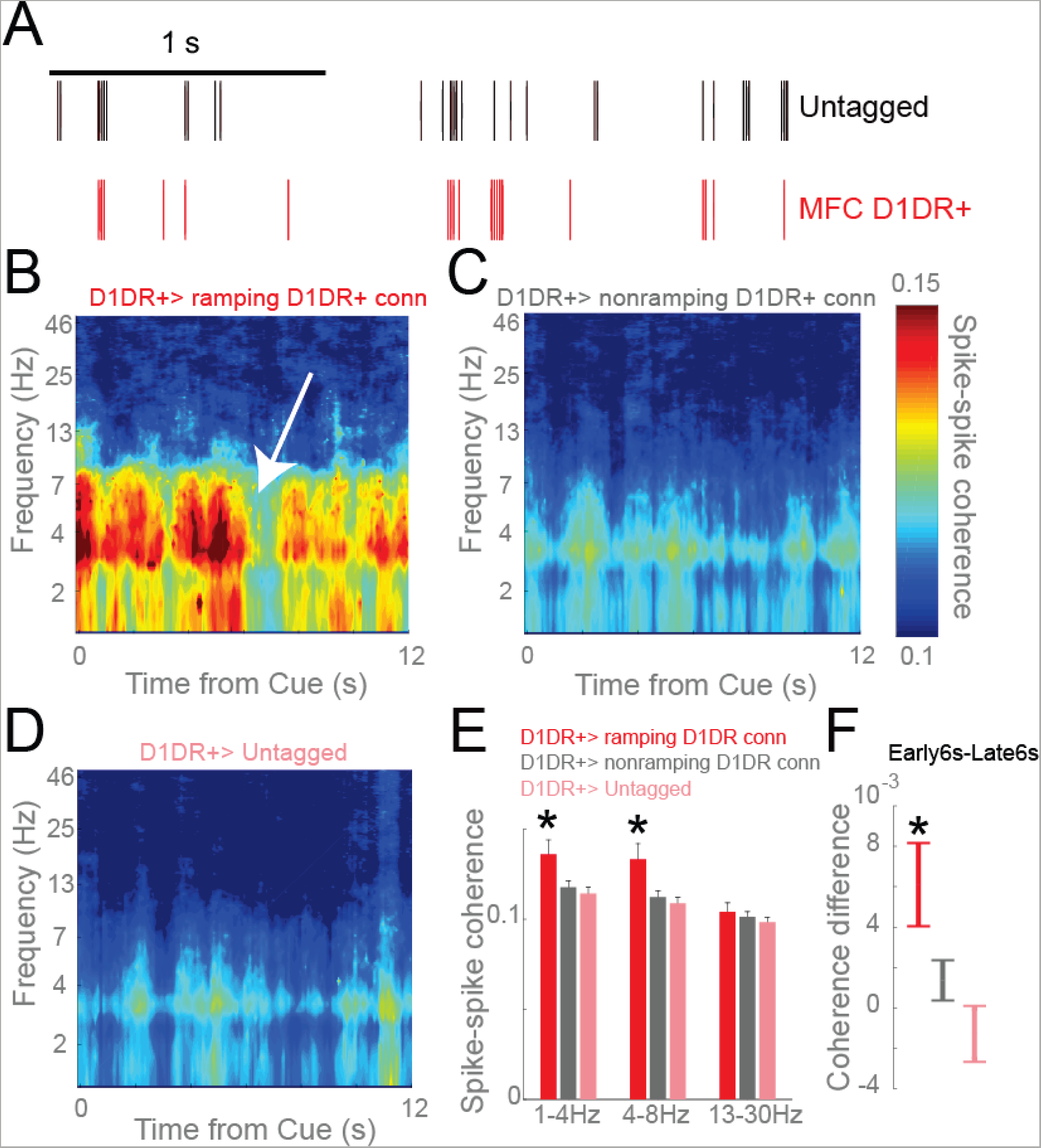
MFC D1DR+ neurons have delta/theta coherence with connected ramping neurons. A) We examined spike-spike coherence, which involves synchrony between the spike trains of two neurons. B) MFC D1DR+ neurons had marked cue-triggered delta/theta coherence that was prominent early in the interval (white arrow appears to be a point of transition). This pattern was not apparent with non-ramping connected neurons or with untagged MFC neurons. E) In addition, MFC D1DR+ neurons had stronger coherence with connected ramping neurons than with non-ramping or untagged neurons. F) MFC D1DR+ neurons had significantly more coherence early in the interval compared with late in the interval (see white arrow in (B), where a transition at ~6 s is apparent). Data from 4679 combinations of 93 D1DR+ neurons, 149 D1DR+ Connected neurons, and 72 untagged MFC neurons in 6 mice; *=p<0.05.

To test the behavioral significance of these interactions, we optogenetically stimulated MFC D1DR+ neurons at delta frequencies (2 Hz) early vs. late in the interval. Our prior work indicates that MFC D1DR+ stimulation for the entire 12-s interval could compensate for behavioral deficits of dopamine depletion (Kim et al. 2017). Data in Figure 5F predict that stimulation should have different effects in the interval, with the first 6 s being more effective as delta interactions are stronger earlier in the interval. To test this prediction, we stimulated MFC D1DR+ neurons from 0-6 s (early) and from 6-12 s (late) at delta frequencies (2 Hz), which would exogenously drive delta coherence among MFC D1DR+ neurons. We measured interval-timing performance by the ‘curvature’ of time-response histograms during behavior. This measure is based on the cumulative distribution of time-response histograms and is independent of overall response rate. Values range between 0 and 1, with impaired performance leading to flatter curvature values closer to 0 (Fry et al. 1960; Narayanan et al. 2012b; Parker, Chen, et al. 2014; Kim et al. 2017). Consistent with our previous studies, we found no effects in mice with intact mesocortical dopamine circuits (Fig 6A). However, these mice are likely at maximal performance, thus limiting this experiment by a ceiling effect (Vijayraghavan et al. 2007; Cools and D’Esposito 2011; Narayanan, Rodnitzky, et al. 2013; Kim et al. 2017). To explore if MFC D1DR+ stimulation was effective in animals with interval-timing deficits, we depleted dopamine in the VTA using the neurotoxin 6-OHDA, which impairs interval timing (VTA-Saline: 0.19 ± 0.06 vs VTA-6-OHDA: 0.07 ± 0.08; t_(11)_=3.3, p<0.01). For these VTA-6OHDA animals there were differential effects of 2-Hz stimulation early vs. late in the interval (t_(5)_=6.2, p<0.002; Fig 6B). As in our previous study with whole-interval stimulation (Kim et al. 2017), stimulation early in the interval was sufficient to improve interval timing and rescue timing deficits in these animals (early 6s vs. non-stim: t_(17)_=-2.6, p<0.02). No effects were found in control sessions with mCherry in D1-Cre mice (Fig 6C). These data support the idea that cue-triggered delta coherence between D1DR+ neurons and ramping neurons early in the interval is key for interval-timing performance.

**Figure 6:**
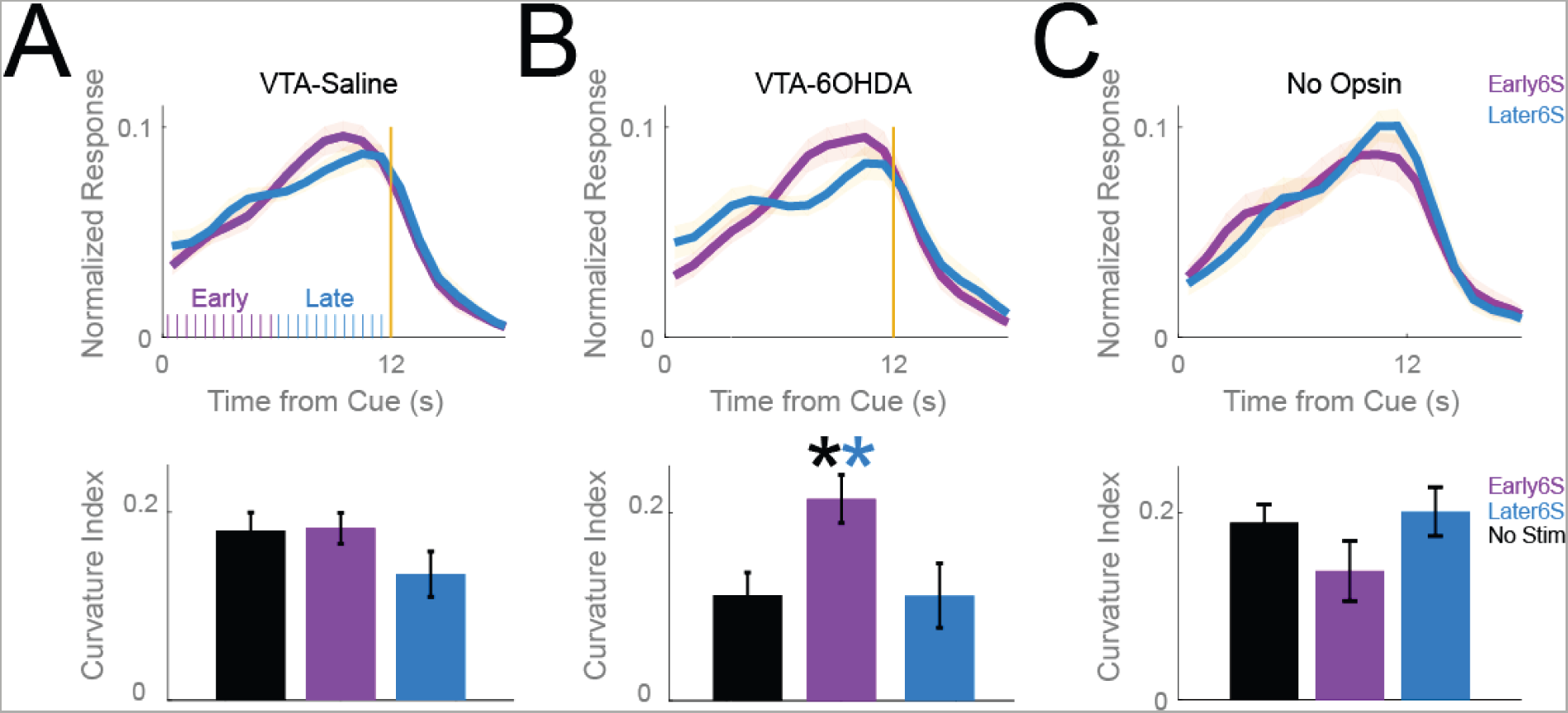
MFC D1DR+ delta stimulation early in the interval can rescue deficits in VTA-6OHDA animals. Time-response histograms for A) VTA-Saline animals (purple ticks indicate laser stimulation pulses early in the interval, blue ticks indicate laser stimulation pulses late in the interval), B), VTA-6OHDA animals, and C) a separate group of 6 control D1-Cre mice expressing virus but no opsin. Blue – MFC D1DR+ 2 Hz stimulation late in the delay from 6-12 s; Purple – stimulation from 0-6 s. Data from 18 mice, *=p<0.05.

## Discussion

We tested the hypothesis that frontal D1DR+ neurons strongly exhibited time-related ramping, a key temporal signal, during interval timing. We recorded from optogenetically-tagged MFC D1DR+ neurons and connected neurons as rodents performed an interval-timing task. While ~23% of MFC D1DR+ neurons exhibited ramping activity, this was less than we observed among other MFC neurons. As such, our data did not provide evidence to support our hypothesis. However, we found that MFC D1DR+ neurons had cue-triggered delta coherence with connected ramping neurons that was strongest early in the interval, predicting that optogenetically increasing delta coherence among these neurons would affect interval-timing performance. In line with this idea, 2-Hz stimulation of MFC D1DR+ neurons early in the interval compensated for behavioral deficits caused by mesocortical dopamine depletion. Our results suggest that MFC D1DR+ neurons provide input to MFC ramping neurons via delta/theta interactions early in the interval. This interaction provides insight into how MFC D1DR+ neurons might support cognitive processing.

Delta/theta frequencies in MFC represent the need for cognitive control (Cavanagh et al. 2012; Narayanan, Cavanagh, et al. 2013; Cavanagh and Frank 2014; Chen et al. 2016). These signals depend on cortical D1DRs and are attenuated in human patients with schizophrenia and Parkinson’s disease (Parker, Chen, et al. 2015; Kim et al. 2017; Parker et al. 2017a) as well as in rodent models (Parker, Chen, et al. 2015; Kim et al. 2017; Parker et al. 2017b). During interval timing, the need for cognitive control is triggered by the cue, which initiates temporal processing (Buhusi and Meck 2005; Meck et al. 2008). In fixed-interval timing tasks, the cue functions as a reward-predictive CS+ involving phasic dopamine release from midbrain dopamine neurons (Schultz 1997; Fonzi et al. 2017). D1DR neurons respond to this phasic dopaminergic release early in the interval and initiate temporal processing via delta-range coherence with MFC ramping neurons.

Our findings go beyond correlative evidence because we show that MFC D1DR+ stimulation improves interval timing (Narayanan et al. 2012b; Kim et al. 2017; Parker et al. 2017b). This is only true when mesocortical dopamine circuits are disrupted, leading to less task-related MFC delta/theta power (Parker, Chen, et al. 2014; Kim et al. 2017). Stimulation is not consistently effective in animals with intact mesocortical dopamine circuits, likely because dopaminergic circuits are close to optimal network function (Vijayraghavan et al. 2007; Cools and D’Esposito 2011; Narayanan, Rodnitzky, et al. 2013; Kim et al. 2017). Our previous stimulation protocols delivered delta stimulation of D1DR+ neurons at a consistent phase during the interval (Narayanan et al. 2012b; Kim et al. 2017; Parker et al. 2017b). If MFC D1DR+ neurons exhibited time-related ramping, this stimulation would have further disrupted interval timing by replacing a time-dependent signal (ramping) with a time-independent signal (a constant 2-Hz firing rate over the interval). We found that constant stimulation of MFC D1DR+ neurons both created time-related ramping among untagged MFC neurons and improved interval-timing behavior (Kim et al. 2017). Findings in Fig 5B&F indicate that delta coherence phase-locked to the interval is sufficient for the effects of MFC D1DR+ neuron stimulation on behavior. These data support a role for oscillatory activity in temporal encoding, as predicted by oscillatory neuronal network models of timing (Matell and Meck 2004; Meck et al. 2008). These models predict phase-locking at response; by contrast, we find phase-locking at the cue. Delta coherence in Figure 5 implies that oscillatory structure early in temporal intervals might be a meaningful temporal signal among MFC neurons. These interactions may help initiate drift-diffusion dynamics that encode time (Simen et al. 2011) via ramping or other mechanisms (Matell and Meck 2004; Latimer et al. 2015; Narayanan 2016)

D1DRs play a critical role in working memory in lateral prefrontal regions in primates (Sawaguchi and Goldman-Rakic 1991, 1994; Williams and Goldman-Rakic 1995; Goldman-Rakic et al. 2004), which are distinct from rodent MFC (Preuss 1995; Narayanan and Laubach 2017). Our group has shown that MFC D1DR+ is critically required for timing behaviors (Narayanan et al. 2012b; Parker, Alberico, et al. 2013) and time-related ramping among MFC neurons (Parker, Narayanan, et al. 2014; Parker, Ruggiero, et al. 2015), but it remains to be seen how these insights relate to other tasks involving cognitive control and prefrontal brain areas in other species.

Our study is limited in making anatomical inferences because we cannot directly visualize recorded neurons and their connectivity. Prefrontal neurons are densely and recurrently connected via local microcircuits and distant projections (Constantinidis et al. 2001; Han et al. 2017). Because of technical limitations, we cannot specify the exact pathway by which MFC D1DR+ neurons entrain ramping neurons or the synaptic connectivity patterns of MFC D1DR+ neurons to other MFC neurons. Such insights likely require correlation of optogenetic approaches with techniques that have better spatial resolution, such as two-photon imaging. It is not clear, however, if these techniques can detect delta/theta coherence (Chen et al. 2013). These data provide novel insight into how dopamine influences cognitive processing in frontal circuits, which could have relevance for disorders such as Parkinson’s disease and schizophrenia.

## Methods

### Transgenic Mice

These experiments used identical procedures to our prior work (Kim et al. 2017). Briefly, we used mice in which Cre-recombinase was driven by the D1DR receptor promoter (*Drd1a-cre*^+^; derived from Gensat strain EY262; aged 3 months; 25-32 g), or littermate controls. Mice were bred and verified by genotyping using primers for D1-Cre recombinase transgene (D1-Cre-F: AGG GGC TGG GTG GTG AGT GAT TG, D1-Cre-R: CGC CGC ATA ACC AGT GAA ACAGC). Mice consumed 1.5-2 g of food pellets (F0071, BioServ) during each behavioral session and additional food was provided 1-3 hours after each behavioral session in the home cage. Single housing and a 12 hour light/dark cycle were used. All experiments took place during the light cycle. Mice were maintained at ~85-90% of their free-access body weight during the course of these experiments for motivation. All procedures were approved by the Animal Care and Use Committee at the University of Iowa #4071105. A total of 12 mice were used for recording experiments (6 D1-Cre+ control mice for recording experiments with saline injected into the VTA and 6 D1-Cre+ mice with dopamine depletion). Eighteen separate mice were used for stimulation experiments: 6 control D1-Cre+ mice expressing ChR2 in the MFC with saline injected into the VTA, 6 D1-Cre+ mice expressing ChR2 in the MFC with mesocortical depletion, 6 control D1-Cre+ mice expressing control virus in the MFC.

Mice were trained to perform an interval-timing task with a 12 s interval according to methods described in detail previously [26]. Time-response histograms were normalized to total responses to investigate timing, independent of response rate. We quantified interval-timing performance by measuring the curvature of time-response histograms by calculating the deviation of the cumulative sum from a straight line; curvature has been used for over 50 years and in our past work to quantify interval timing behavior (Fry et al. 1960; Narayanan et al. 2012a; Parker, Chen, et al. 2014; Emmons et al. 2017; Kim et al. 2017; Parker et al. 2017a). Curvature indices are higher with more ‘curved’ time-response histograms. As above, all behavioral data was tested for normality.

Mice trained in the 12 s interval-timing task were implanted with 16-channel 50 micron stainless-steel recording electrodes and an optical fiber in the MFC (Microprobes) [26]. Surgical procedures, neurophysiological recordings, neuronal analyses, and time-frequency analyses of mouse LFPs were conducted identical to methods described in detail previously (Emmons et al. 2016; Kim et al. 2017; Parker et al. 2017a)

### Optogenetics

We used an AAV construct with floxed-inverted channelrhodopsin along with mCherry (UNC Viral Core; AAV5-EF1a-DIO-hChR2(H134R)-mCherry; AAV-DIO-ChR2) (Cardin et al. 2009). Control virus expressing mCherry. When delivered to transgenic D1-Cre+ mice, Cre recombination leads to high expression driven by an EF-1a promoter selectively in neurons expressing D1DRs. Mice were injected with AAV-DIO-ChR2 into the prefrontal cortex (Mouse: AP: +1.8, ML −0.5, DV −1.5), with immediate placement of an optical fiber cannula (200 µm core, 0.22NA, Doric Lenses). The injection consisted of 0.5 µL of approximately 1-8x10^12 infectious particles per milliliter.

On testing days, D1-Cre+ mice with optical cannula were connected to the optical patch cable through a Zirconia ferrule (Doric Lenses) without anesthesia. Light was generated from a 473 nm DPSS laser source (OEM Laser Systems) and an optical rotary joint (Doric Lenses) was used to facilitate animal rotation during performance of the interval-timing task. During testing, each mouse performed the fixed-interval timing task for 1 hr with light delivered with specific frequencies of stimulation. Specific frequencies of laser light were generated by TTL signals sent through a microcontroller controlled by the operant behavior computer. In stimulation sessions, light was delivered from 0 to 6 s or 6 to 12 s during the fixed-interval at 0 and 2 Hz with a pulse width of 5 ms during randomly selected trials (33% for each condition – early, late, and 0 Hz; 0 Hz meant that the laser was off and no laser light was delivered). The power output of laser was adjusted to be 8 mW at the fiber tip before every experiment, power measurements verified that the laser reached 90% power within 0.74 ms of TTL triggers and maintained 8 mW with <5% error.

### Neuronal Ensemble Recordings

Neuronal ensemble recordings in the MFC were made using a multi-electrode recording system (Plexon). Raw signal was amplified with total gain of 5000 and high-pass filtered at 0.05Hz and recorded with 16bit resolution at 40k Hz sampling rate. To detect spikes, raw signals were re-referenced using common median referencing to minimize potential non-neural electrical noises and band-passed filtered between 300 and 6000 Hz offline. Spikes were detected with a threshold of 5 median absolute deviations. Plexon Offline Sorter was used to sort single units and to remove artifacts. PCA and waveform shape were used for spike sorting. Spike activity was analyzed for all cells that fired at rates above 0.1 Hz. Statistical summaries were based on all remaining neurons. No subpopulations were selected or filtered out of the neuron database. LFP was recorded with bandpass filters between 0.05 and 1000 Hz. Analysis of neuronal activity and quantitative analysis of basic firing properties were carried out with custom routines for MATLAB. All behavioral events and laser triggers were recorded simultaneously using TTL inputs. Peri-event rasters and average histograms were constructed around trial start, and laser light pulse.

Spike-spike coherence was calculated between pairs of neurons recorded during the same sessions. The magnitude of the coherence for trial-aligned spike trains was calculated using the Chronux toolbox with multi-taper Fourier analysis (Mitra and Bokil 2008). Calculations were performed using following parameters: window size=1s; moving step=0.1s; number of tapers=5. To compare across neurons with different firing rates, spike-rate differences are adjusted by estimating a correction factor that is conceptually equivalent to spike bootstrapping procedures (Aoi et al. 2015).

### Histology

When experiments were complete, mice were anesthetized and sacrificed by injections of 100 mg/kg sodium pentobarbital. All mice were intracardially perfused with 4% paraformaldehyde. The brain was removed and post-fixed in paraformaldehyde overnight, and immersed in 30% sucrose until the brains sank. 50µm sections were made on cryostat (Leica) and stored in PBS. Standard immunostaining procedures were performed in free-floating brain sections. Primary antibodies to Cre (mouse anti-Cre; Millipore-MAB 3120; 1:500), D1 receptor (rat anti-D1 dopamine receptor; Sigma-D2944; 1:200), tyrosine hydroxylase (rabbit anti-TH; Millpore-AB152; 1:500) were incubated overnight at 4 ºC. Sections were visualized with Alexa Flour fluorescent secondary antibodies (goat anti-mouse IgG Alexa 633, goat anti-rat IgG Alexa 568, and goat anti-rabbit IgG Alexa 488; ThermoFisher; 1:1000) matched with the host primary by incubating for 2 hours at room temperature. Images were captured on Leica SP5 laser scanning confocal microscope or Zeiss Apotome.2 Axio Imager.

## Acknowledgements

This work was funded by The National Institute of Neurological Disorders and Stroke R01NS078100/K08 NS078100, The National Institute of Mental Health, NARSAD Young Investigator Grant from Brain & Behavior Foundation, and grant #2014/22817-1. We’d like to thank Eric Emmons and Ben De Corte for feedback on this manuscript, and Sangwoo Han for technical assistance.

